# Capturing the Diversity of Subsurface Microbiota – Choice of Carbon Source for Microcosm Enrichment and Isolation of Groundwater Bacteria

**DOI:** 10.1101/517854

**Authors:** Xiaoqin Wu, Sarah Spencer, Eric J. Alm, Jana Voriskova, Romy Chakraborty

## Abstract

Improved and innovative enrichment/isolation techniques that yield to relevant isolates representing the true diversity of environmental microbial communities would significantly advance exploring the physiology of ecologically important taxa in ecosystems. Traditionally, either simple organic carbon (C) or yeast extract is used as C source in culture medium for microbial enrichment/isolation in laboratory. In natural environment, however, microbial population and evolution are greatly influenced by the property and composition of natural organic C. In this study, 8 types of organic C sources were fed to intrinsic groundwater microbes collected at Oak Ridge Reservation Field Research Center (ORR-FRC) background site for a 30-day incubation period to investigate the response of indigenous bacterial communities to different C sources. The tested C sources included simple organic C (glucose, acetate, benzoate, oleic acid, and cellulose) that are either traditionally used as C source in bacterial culture medium or present in natural environments; naturally occurring undefined complex C (bacterial cell lysate and sediment-derived natural organic matter (NOM)); as well as vitamin mixture which is a commonly used ingredient in culture medium. Our results clearly indicate that natural complex C substrates served better in enriching diverse bacteria compared to other C sources. Microcosms amended with small organic C (glucose, acetate, benzoate, or oleic acid) showed significantly lower biodiversity than control groups, dominated by only a few phyla of bacteria such as *Proteobacteria* and *Bacteroidetes* which are commonly isolated and already have diverse representative isolates, while those amended with natural complex C (cell lysate or NOM) displayed significantly higher biodiversity than control groups, in which three phyla (*Verrucomicrobia, Planctomycetes*, and *Armatimonadetes*) that are poorly represented in published culture collections were abundantly enriched. Further isolation of pure bacterial strains from complex C-amended enrichments led to 51 species representing 4 phyla, 13 orders. Furthermore, 5 isolates with low similarities to published strains were considered to be novel. Results from this study will aid in the design of better cultivation and isolation strategy for maximize the diversity of organisms recovered from subsurface environment.

## Introduction

Using 16S ribosomal RNA (rRNA) gene or metagenomics surveys from a wide range of habitats, scientists have uncovered an astounding diversity of bacteria living on our planet. Yet, only a small portion (<1%) of bacteria on Earth have been successfully cultivated^1, 2^ and about half of those reported bacterial phyla still lack cultivated representatives^3^. While rapid technological advances are being made in developing modern molecular tools such as metagenomics, metaproteomics, and metatranscriptomics to identify key microbial species and metabolic potential in a given environment, the complete interpretation of the data is constrained by the unavailability of reference genomes and isolates to serve as reference data, and validate the hypotheses that emerge from powerful omics-based data^4^.

For years scientists have been trying to develop cultivation/isolation methods and techniques, such as modification of growth media/conditions, use of diluted medium or serial dilution culture^5, 6^, iChip^7^, diffusion chamber^8-10^, etc., to cultivate diverse environmental bacteria especially those ‘unculturable’ species under laboratory conditions^11^. Successful cultivation of bacteria critically depend on the choice of appropriate growth media and incubation conditions^12^. For most chemoheterotrophs, organic carbon (C) source is a key ingredient in culture medium since C incorporates into cellular matter for bacterial growth and serves as electron donor for energy transfer in bacteria. Traditionally, yeast extract or simple organic compounds, e.g., glucose, acetate, lactate, pyruvate, and casamino acids, are added as C source either individually or as a mixture to the cultivation medium^13^. However, these labile C usually lead to selective and biased growth of only specific microbes^14, 15^.

To increase the diversity of enrichment/isolates from environmental samples, researchers have utilized media that mimic environmental habitats of microbes, and found that some previously uncultivable microbes could be grown in pure culture if provided with chemical components that mimics their natural environments^8, 16-18^. Natural organic matter (NOM) is the naturally occurring heterogeneous organic C source for most microbes in natural ecosystems, consisting of complex C that differ in molecular mass, solubility, structure, and functionality^19-21^. Recently, Nguyen et al. applied soil-extracted NOM as an ingredient in culture medium and obtained diverse bacterial isolates including those previously uncultured and novel species from soil^22^.

To date our knowledge and understanding of microbial ecology in the subsurface are still extremely scarce. Of the published 16S rRNA gene sequences in public databases, only <8% are derived from subsurface organisms, and only a small fraction of those are represented by genomes or isolates^23^. The lack of innovation and alternative to cultivation has severely limited the ability of microbiologists to characterize microbes that catalyze key biogeochemical processes in subsurface ecosystems. In this study, we aimed to explore the influence of naturally occurring complex C on cultivation and isolation of bacteria from subsurface groundwater from the Field Research Center (FRC) in Oak Ridge, TN. We employed microcosm enrichments and fed microbes from groundwater with bacterial cell lysate or sediment-extracted NOM as C source. As a comparison, we also included enrichment groups that were amended with relatively simple C sources, i.e., small sugar (glucose), small metabolites (acetate or benzoate), fatty acid (oleic acid), polysaccharide (cellulose), or mixed vitamins. We included mixed vitamins since they are usually added as supplements to bacterial growth media^24, 25^. This knowledge would benefit for optimizing the strategy for cultivation/isolation of relevant isolates representing the microbial diversity in the subsurface, which is critical for understanding the physiology of ecologically important taxa in subsurface ecosystems.

## Materials and Methods

### Preparation of C stock solutions

Standards of glucose, sodium acetate, sodium benzoate, cellulose, oleic acid, vitamins, and thioctic acid were purchased from Sigma-Aldrich (St. Louis, MO). Stock solutions of glucose, sodium acetate, and sodium benzoate were prepared by dissolving the chemical in MilliQ-water (18.2 MΩ·cm, 0.22 μm membrane filtered) at 200 mM, 200 mM, and 50 mM, respectively, followed by filter-sterilization with filtration system (0.22 μm pore-sized, polyethersulfone (PES), Corning). Oleic acid and cellulose were added to MilliQ-water at an initial concentration of 50 g/L and 20 g/L, respectively, followed by autoclave sterilization. Since oleic acid and cellulose are generally insoluble, their concentrations in water are expressed as initial grams per liter. Stock solution of mixed vitamins, including vitamin B_1_, B_2_, B_3_, B_5_, B_6_, B_7_, B_9_, B_10_, B_12_, and thioctic acid, was prepared in MilliQ-water according to the recipe reported by Balch et al.^24^ (Supporting Information), and then filter-sterilized (0.22 μm pore-sized, PES, Corning).

Preparation procedure of cell lysate stock solution was modified based on published methods^26, 27^. A strain of *Pseudomonas fluorescens*, which was previously isolated in our lab from groundwater collected at Oak Ridge FRC background site was grown in Luria broth (LB) liquid medium at 30 °C under aerobic condition until the optical density at 600 nm (OD_600_) reached stationary phase. A 30-ml aliquot of the culture was harvested and centrifuged at 6,000 *g* for 20 min. The supernatant was removed and the pellet was washed by MilliQ-water for three times and re-suspended in 10 ml of MilliQ-water. A two-step lysis procedure was applied, including autoclaving and sonication in water bath for 2 hrs. Then the solution was centrifugation at 6,000 *g* for 20 min. The supernatant was decanted and filtered through a syringe filter (0.2 μm pore-sized, PES, Thermo Scientific). The filtrate was stored at 4 °C until use. Total organic C (TOC) content of the filtrate, i.e., cell lysate stock solution, was 2.67 g/L, measured by TOC-5050A Total Organic Carbon Analyzer (Shimadzu, Japan).

The sediment sample for NOM extraction was collected from a background well FW305 at ORR-FRC, at the depth of 0.3-1.1 m below ground surface. The potential bioavailable fraction of sediment NOM was extracted according to the method previously developed in our lab^28^. Briefly, the freeze-dried sediment sample was extracted with Milli-Q water via rotary shaking (170 rpm) overnight at 35 °C, and then sonicated in water bath for 2 hrs. The ratio of water and sediment was 4:1 (w/w). After extraction, the water-sediment mixtures were centrifuged at 6000 *g* for 20 min. The supernatant was decanted and sterilized using filtration system (0.22 μm pore-sized, PES, Corning). The filtrate containing water-extractable NOM was freeze-dried, and the lyophilized material was stored at -20 °C until use.

### Microcosm enrichment

Groundwater sample was collected from a background well GW305 adjoining to the sediment well FW305 at ORR-FRC in April, 2016. After being collected, the groundwater was shipped immediately to the lab with ice packs, and stored at 4 °C for up to 1 week. At the time of sampling, the groundwater temperature was measured to be 15.4 °C, pH was 6.37, dissolved oxygen (DO) was 1.39 mg/L, and TOC was 5.9 mg/L. The DO in groundwater exceeded 0.5 mg/L, indicating the groundwater sample’s redox state was oxic (Ohio EPA, http://epa.ohio.gov/Portals/28/documents/gwqcp/redox_ts.pdf).

Microcosm experiments were performed in pre-sterilized 250 ml-flasks, each containing 89 ml of filtered groundwater (0.22 µm pore-sized, PES, Corning) as culture medium, 10 ml of unfiltered groundwater (cell density: 2.1 × 10^6^ cells/ml) as inoculum, and 1 ml of C stock solution. For oleic acid and cellulose amended groups, the C stock solutions were shaken thoroughly to mix the solution and undissolved chemicals well before adding to the culture. For sediment NOM amended group, the lyophilized material was fully dissolved in filtered groundwater at 200 mg/L, and then the solution was filter-sterilized (0.22 µm pore-sized, PES, Corning). TOC content of the filtrate was measured to be 48.4 mg/L. A 90 ml aliquot of the filtrate (containing sediment NOM) was added with 10 ml of unfiltered groundwater to form a microcosm.

A control group without any C amendment was included in this study, containing 90 ml of filtered groundwater and 10 ml of unfiltered groundwater. All groups were performed in six replicates, and 1 blank control (without inoculum) was included in each group to monitor potential microbial contamination during incubation. All microcosms were incubated aerobically at 25 °C in the dark for up to 30 days, with rotary shaking at 100 rpm. At each sampling time point (day 10, 20, and 30), a 10 ml aliquot of subculture was sampled using a volumetric pipette. Microbes were collected by filtration through a membrane filter (0.2 µm pore-sized, PES, 25 mm, Sterlitech Corp.). The filter was then removed from the syringe filter holder and kept frozen at -80 °C until DNA extraction.

### DNA extraction for microbial community analysis

Before performing DNA extraction, the filters were cut into 2 mm-wide stripes using sterile blades and put into DNA extraction tubes provided in PowerMax Soil DNA Isolation Kit (MO BIO Laboratories, Inc., Carlsbad, CA). DNA was extracted following the manufacturer’s protocol, and quantified using the Qubit dsDNA HS Assay Kit (Life Technologies, Eugene, OR) with a Qubit fluorometer (Invitrogen, Eugene, OR). The extracted DNA samples were stored at -20 °C until further processing.

### 16S rRNA gene amplicon library preparation

We completed a two-step PCR protocol to first amplify the 16S rRNA gene V4 variable region, then add Illumina barcodes and adapters for sequencing. The DNA samples were each aliquot into one of three randomized plate layouts in a laminar flow hood. Up to 25 μl of each sample was transferred, with eight wells per plate left open for amplification negative controls.

Before the first step PCR, all samples were subjected to a qPCR at multiple dilutions to determine target dilutions and threshold cycles for the first step. We used 16S rRNA gene primers PE16S_V4_U515_F and PE16S_V4_E786R (Supplementary Table S1). Both 1:1 and 1:10 dilutions of each sample were prepared in duplicate with 0.5X SYBR Green I nucleic acid gel stain (Sigma-Aldrich, St. Louis, MO), plus 280 nM each primer and the standard reagents in the Phusion High-Fidelity PCR Kit (New England BioLabs, Ipswich, MA). Samples were then cycled under the following qPCR conditions: 98 °C 30 sec; 30 cycles of 98 °C 30 sec, 52 °C 30 sec, 72 °C 30 sec; 4 °C hold. Threshold cycles were calculated and dilutions were prepared to normalize samples and ensure consistent amplification cycles across plates. PCR under the same conditions, minus the SYBR Green, was completed in quadruplicate for each sample, then quadruplicate sets were pooled and purified with Agencourt AMPure XP Beads according to the manufacturer’s protocol (Beckman Coulter, Brea, CA).

The second step PCR was used to add sample indices and final Illumina adaptors to the 16S rRNA gene amplicons. Reactions were compiled using the Phusion High-Fidelity PCR Kit according to the manufacturer’s instructions, with 420 nM indexing primers PE-III-PCR-F and PE-IV-PCR-R (Supplementary Table S1), then cycled under the following conditions: 98 °C 30 sec; 7 cycles of 98 °C 30 sec, 83 °C 30 sec, 72 °C 30 sec; 4 °C hold. Final libraries were purified with Agencourt AMPure XP Beads according to the manufacturer’s protocol, then quantified and pooled prior to 2 × 250 paired-end sequencing on an Illumina MiSeq. Data are available on the SRA under accession.

### 16S rRNA gene amplicon data processing and operational taxonomic unit (OTU) analysis

Raw reads were quality filtered and clustered into operational taxonomic units (OTUs) primarily with the QIIME software package^29^ using default parameters unless otherwise noted. Paired-end reads were joined with the join_paired_ends.py command, then barcodes were extracted from the successfully joined reads with the extract_barcodes.py command (and additional parameters -c barcode_in_label, -l 16, -s ‘#’). Quality filtering was accomplished with split_libraries_fastq.py (--barcode_type 16, --min_per_read_length_fraction 0.40, -q 20, --max_barcode_errors 0, -- max_bad_run_length 0, --phred_offset 33). We checked for the correct forward and reverse primers with a custom script and exported reads with primers removed and length trimmed to 225 bp. Finally, chimeric sequences were removed using identify_chimeric_seqs.py (-m usearch61, --suppress_usearch61_ref) followed by filter_fasta.py.

After quality filtering, reads were clustered into 97% OTUs, classified against a 16S rRNA database, and aligned in order to build phylogenetic trees. We ran the QIIME commands pick_otus.py, pick_rep_set.py (-m most_abundant), and make_otu_table.py to produce the OTU table. The RDP classifier was used to assign taxonomy with default parameters and the 16S rRNA training set 16^30^. Representative sequences from OTUs with > 0.1% and > 5% abundance in at least one experimental sample were selected for alignment and tree construction. Alignment was completed with SINA 1.2.11 using the SILVA reference alignment (SSURef_NR99_128_SILVA_07_09_16_opt.arb)^31^. Trees were constructed FastTree 2.1.9 with a generalized time-reversible model^32^.

### Bacterial isolation

The bacterial cell lysate-amended and NOM-amended enrichments at each time point were used as inoculums for further isolation. Each selected enrichment sample was streaked on complex C agar plate, which was prepared using the same medium as corresponding liquid enrichment with 1.5% agar (BD Biosciences, USA). We also streaked each selected enrichment sample on diluted culture media (with 1.5% agar), i.e., 1/25 LB, 1/25 tryptic soy broth (TSB), and 1/10 Reasoner’s 2A (R2A), to obtain as many colonies as possible. The plates were incubated at 27 °C in the dark. Bacterial colonies were repeatedly streaked until single colonies were obtained. The single colony was picked from the plate and transferred to 3 ml of corresponding liquid medium. The liquid cultures were incubated at 27 °C in the dark for up to 3 weeks before DNA extraction.

### Species identification

Genomic DNA of isolates were extracted using a PureLink Genomic DNA Mini Kit (Invitrogen, United States) following manufacturer’s protocol. 16S rRNA genes were amplified (initial denaturation step at 98°C for 5 min, followed by 30 cycles at 95°C for 30 s, 50°C for 30 s and 72°C for 2 min, followed by a final step at 72°C for 3 min) using the eubacterial primers 27F (AGA GTT TGA TCC TGG CTC AG) and 1492R (ACG GCT ACC TTG TTA CGA CTT) purchased from Integrated DNA Technologies, Inc. (USA). Cleanup of PCR products and DNA sequencing were performed at University of California Berkeley DNA Sequencing Facility. The PCR products were sequenced using the internal primers 27F and 1492R. Sequences were obtained by Sanger sequencing with ABI 3730XL DNA Analyzers (ThermoFisher, United States). Consensus sequences (1200–1400 base pairs) from forward and reverse sequences were generated using Geneious (version 9.1.3). A subset of isolate’s sequences was deposited in Genbank under the access codes XXX to XXX (Table 1). For bacterial isolates identification, Megablast (Genbank) BLAST was used to obtain the top hits.

**Table 1.** Isolates from microcosm enrichments amended with cell lysate (CL) or sediment NOM.

| Phylum | Order | Isolate ID | Accession number | Highest 16S rRNA gene sequence similarity [GenBank sequence ID] | Similarity | C source in the inoculum |
| --- | --- | --- | --- | --- | --- | --- |
| *Actinobacteria* | *Micrococcales* | FW305-C-20 |  | *Microbacterium maritipicum* strain DSM 12512 [NR_114986.1] | 99 | CL |
|  |  | FW305-C-125 |  | *Herbiconiux solani* strain K 134/01 [NR_116995.1] | 99 | NOM |
|  |  | FW305-C-176A |  | *Leucobacter iarius* strain 40 [NR_042414.1] | 99 | CL |
|  |  | FW305-C-184C |  | *Glaciihabitans tibetensis* strain MP203 [NR_133754.1] | 98 | CL |
| *Bacteroidetes* | *Chitinophagales* | FW305-C-49 |  | *Sediminibacterium goheungense* strain HME7863 [NR_133854.1] | 99 | CL |
|  |  | FW305-C-178 |  | *Sediminibacterium salmoneum* strain NJ-44 [NR_044197.1] | 99 | CL |
|  |  | FW305-C-185 |  | *Terrimonas soli* strain FL-8 [NR_159891.1] | 98 | CL |
|  | *Cytophagales* | FW305-C-70 |  | *Flectobacillus roseus* strain GFA-11 [NR_116312.1] | 99 | NOM |
|  |  | FW305-C-80 |  | *Emticicia ginsengisoli* strain Gsoil 085 [NR_041373.1] | 99 | NOM |
|  |  | FW305-C-84 |  | *Dyadobacter sediminis* strain Z12 [NR_134722.1] | 97 | NOM |
|  | *Sphingobacteriales* | FW305-C-75 |  | *Pedobacter ginsengisoli* strain Gsoil 104 [NR_041374.1] | 99 | NOM |
|  |  | FW305-C-21 |  | *Solitalea canadensis* strain DSM 3403 [NR_074099.1] | 86 | CL |
| *Firmicutes* | *Bacillales* | FW305-C-191 |  | *Bacillus firmus* strain NBRC 15306 [NR_112635.1] | 99 | CL, NOM |
|  |  | FW305-C-48 |  | *Brevibacillus agri* strain DSM 6348 [NR_040983.1] | 99 | CL |
|  |  | FW305-C-1 |  | *Brevibacillus nitrificans* strain DA2 [NR_112926.1] | 100 | CL, NOM |
|  |  | FW305-C-190 |  | *Paenibacillus lautus* strain NBRC 15380 [NR_112724.1] | 98 | CL |
|  |  | FW305-C-202 |  | *Paenibacillus naphthalenovorans* strain PR-N1 [NR_028817.1] | 99 | CL |
| *Proteobacteria*<br>(Alpha) | *Caulobacterales* | FW305-C-18 |  | *Brevundimonas vesicularis* strain NBRC 12165 [NR_113586.1] | 100 | CL |
|  |  | FW305-C-128 |  | *Caulobacter fusiformis* strain ATCC 15257 [NR_025320.1] | 99 | NOM |
|  |  | FW305-C-130 |  | *Caulobacter profundus* strain DS48-5-2 [NR_133716.1] | 99 | NOM |
|  | *Rhizobiales* | FW305-C-122 |  | *Afipia broomeae* strain F186 [NR_029200.1] | 99 | NOM |
|  |  | FW305-C-52 |  | *Aminobacter niigataensis* strain DSM 7050 [NR_025302.1] | 99 | CL |
|  |  | FW305-C-92 |  | *Bosea lupini* strain R-45681 [NR_108514.1] | 99 | NOM |
|  |  | FW305-C-101 |  | *Bosea robiniae* strain R-46070 [NR_108516.1] | 99 | NOM |
|  |  | FW305-C-74 |  | *Bosea vestrisii* strain 34635 [NR_028799.1] | 99 | NOM |
|  |  | FW305-C-47 |  | *Devosia insulae* strain DS-56 [NR_044036.1] | 99 | CL, NOM |
|  |  | FW305-C-198 |  | *Methylobacterium oryzae* strain CBMB20 [NR_043104.1] | 99 | NOM |
|  |  | FW305-C-112 |  | *Nordella oligomobilis* strain N21 [NR_114615.1] | 99 | NOM |
|  |  | FW305-C-8 |  | *Rhizobium herbae* strain CCBAU 83011 [NR_117530.1] | 99 | CL |
|  |  | FW305-C-176I |  | *Rhizobium rosettiformans* strain W3 [NR_116445.1] | 96 | CL |
|  |  | FW305-C-134A |  | *Phreatobacter stygius* strain YC6-17 [NR_158009.1] | 99 | NOM |
|  | *Rhodospirillales* | FW305-C-103 |  | *Roseomonas rubra* strain: S5 [NR_152066.1] | 99 | NOM |
|  |  | FW305-C-119 |  | *Roseomonas stagni* strain HS-69 [NR_041660.1] | 98 | NOM |
|  | *Sphingomonadales* | FW305-C-71 |  | *Novosphingobium aquaticum* strain THW-SA1 [NR_148323.1] | 99 | NOM |
|  |  | FW305-C-111 |  | *Novosphingobium aromaticivorans* strain DSM 12444 [NR_074261.1] | 99 | NOM |
|  |  | FW305-C-53 |  | *Novosphingobium subterraneum* strain NBRC 16086 [NR_113838.1] | 99 | CL |
|  |  | FW305-C-94 |  | *Sphingomonas koreensis* strain NBRC 16723 [NR_113868.1] | 99 | NOM |
|  |  | FW305-C-56 |  | *Sphingomonas wittichii* strain RW1 [NR_074268.1] | 99 | CL |
|  |  | FW305-C-54 |  | *Sphingopyxis panaciterrae* strain Gsoil 124 [NR_112561.1] | 99 | CL |
| *Proteobacteria*<br>(Beta) | *Burkholderiales* | FW305-C-28 |  | *Achromobacter deleyi* strain LMG 3458 [NR_152014.1] | 99 | CL |
|  |  | FW305-C-31 |  | *Achromobacter marplatensis* strain B2 [NR_116198.1] | 99 | CL, NOM |
|  |  | FW305-C-25 |  | *Acidovorax wautersii* strain NF 1078 [NR_109656.1] | 99 | CL |
|  |  | FW305-C-176C |  | *Cupriavidus basilensis* strain DSM 11853 [NR_025138.1] | 98 | CL |
|  |  | FW305-C-136 |  | *Curvibacter fontanus* strain AQ9 [NR_112221.1] | 99 | CL, NOM |
|  |  | FW305-C-24 |  | *Pseudacidovorax intermedius* strain CC-21 [NR_044241.1] | 99 | CL, NOM |
|  |  | FW305-C-7 |  | *Variovorax boronicumulans* strain NBRC 103145 [NR_114214.1] | 99 | CL |
|  | *Rhodocyclales* | FW305-C-19 |  | *Dechloromonas agitata* strain CKB [NR_024884.1] | 98 | CL |
| *Proteobacteria*<br>(Gamma) | *Nevskiales* | FW305-C-40 |  | *Hydrocarboniphaga effusa* strain AP103 [NR_029102.1] | 99 | CL |
|  |  | FW305-C-2 |  | *Panacagrimonas perspica* strain Gsoil 142 [NR_112617.1] | 93 | CL, NOM |
|  |  | FW305-C-3 |  | *Sinimarinibacterium flocculans* strain NH6-24 [NR_137419.1] | 96 | CL |
|  | *Pseudomonadales* | FW305-C-5 |  | *Pseudomonas lactis* strain DSM 29167 [NR_156986.1] | 99 | CL |

### Data analysis and statistics

Shannon’s diversity index (*H’*) and multivariate statistics were performed using the R package *vegan*. OTU distributions were transformed into relative abundances using the function *decostand*. These were subjected to Hellinger transformation before calculation of Bray-Curtis dissimilarity matrices comparing community composition between samples. Non-metric multidimensional scaling (NMDS) using function *metaMDS* was performed using these dissimilarity matrices. Multivariate analysis of variance (MANOVA) model was implemented in the *vegan* function *adonis*. Analysis of similarity (ANOSIM) was carried out based on Bray-Curtis dissimilarities in order to evaluate the effect of C amendment and incubation time on community structure.

C utilization was summarized using available replicates for each treatment condition and sampling time point. In this study, an OTU was visualized as actively utilizing a C source if it was present with >1% relative abundance in at least three replicates for any time point (day 10, 20, or 30). We described generalist species as those with all C utilizations (including control group), specialists as those with a single C utilization, and labeled those in between as intermediates. Those without any C designations were either with low abundance in the three time points or showed transient growth between these time points.

## Results

### Complex C sources increased bacterial diversity in microcosm enrichments

In this study, 16S rRNA gene amplicon sequencing resulted in over 10 M prokaryotic 16S rRNA gene reads which were clustered into 3463 OTUs. Only rarefied OTU richness was considered further, in order to compensate for differences in sequencing depth between 144 samples. No DNA was detected in blank controls, suggesting that microbial contamination was negligible during incubation.

Statistical analysis showed that amended C source and incubation time both had significant influences on bacterial community structure (MANOVA/*adonis* and ANOSIM, *p* = 0.001). C source was the major driver of community dissimilarity (MANOVA/*adonis, R*^2^ = 0.56; ANOSIM, *R* = 0.88), whereas incubation time contributed to a lesser extent to the variation (MANOVA/*adonis, R*^2^ = 0.09; ANOSIM, *R* = 0.12). Accordingly, samples were grouped on NMDS ordination diagram based on the type of amended C source. Figure 1 clearly shows that the trajectory of bacterial communities is influenced by C substrates. Bacterial composition in cultures amended with simple small organic C such as glucose, acetate, benzoate, or oleic acid were noticeably similar to each other at the early stage of incubation, and then diversified (Figure 1). In cultures amended with undefined complex C, such as bacterial cell lysate or sediment NOM, the bacterial community separates from other groups early on (Figure 1).

**Figure 1.**
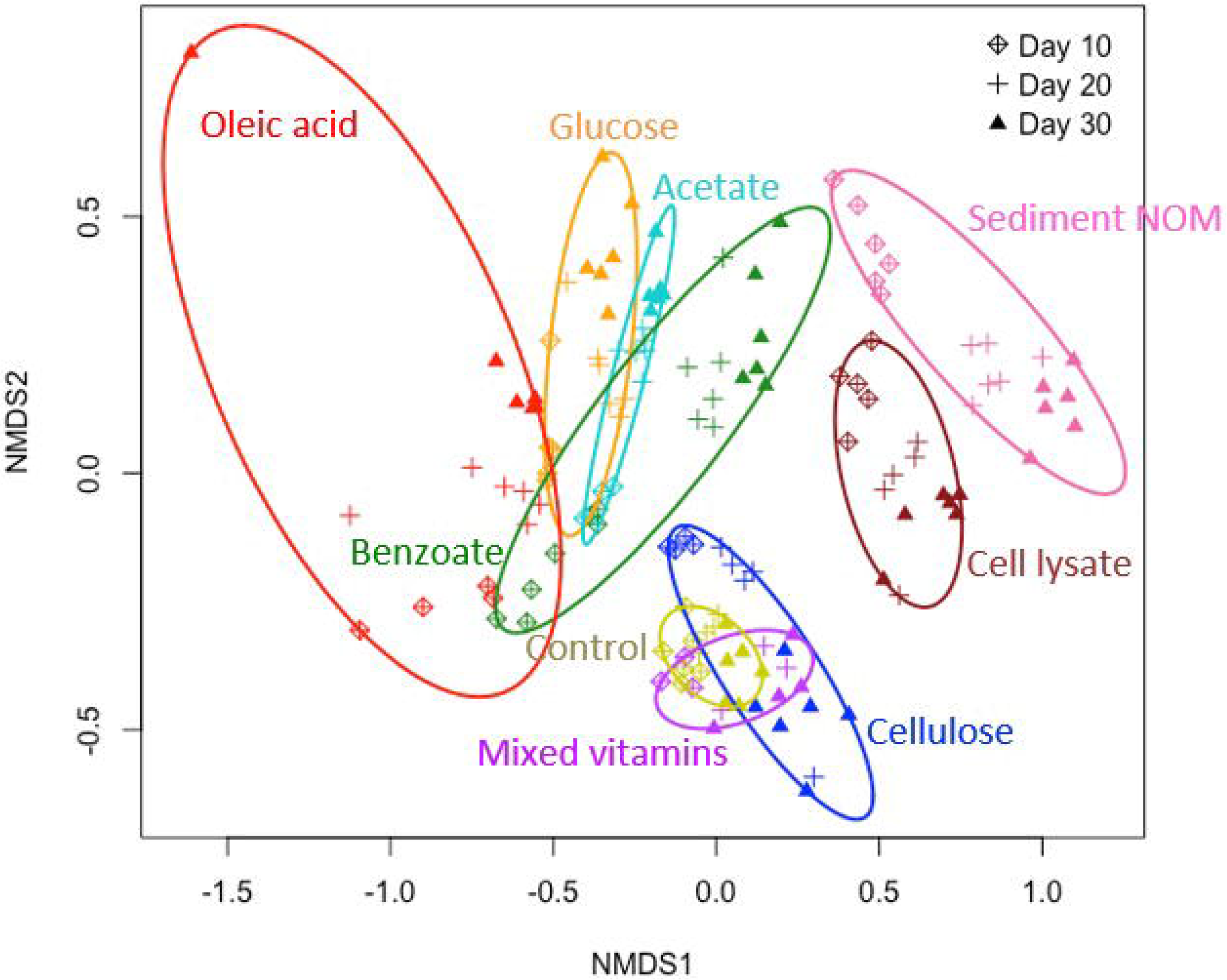
Non-metric multidimensional scaling (NMDS) based on Bray-Curtis dissimilarities of bacterial community composition.

The complexity of C substrates also affected bacterial community diversity, which was evaluated by Shannon’s diversity index (*H’*). As shown in Figure 2, enrichments with simple small organic C (glucose, acetate, benzoate, or oleic acid) have generally lower *H’* values than those in the control group at that corresponding time point, suggesting that simple small organic C sources may decrease community diversity and lead to enrichment of a few bacteria species that preferentially utilize specific C. On the contrary, the *H’* values in enrichments amended with undefined complex C (bacterial cell lysate or sediment NOM) are higher than those in corresponding control group as well as other groups (Figure 2), suggesting that complex C sources encourage cultivation and enrichment of more diverse bacteria compared to simple organic C sources. No significant difference in community composition and diversity was observed between vitamins or cellulose-amended groups and control group (Figure 1 and 2), indicating that these C substrates had insignificant influence on bacterial communities.

**Figure 2.**
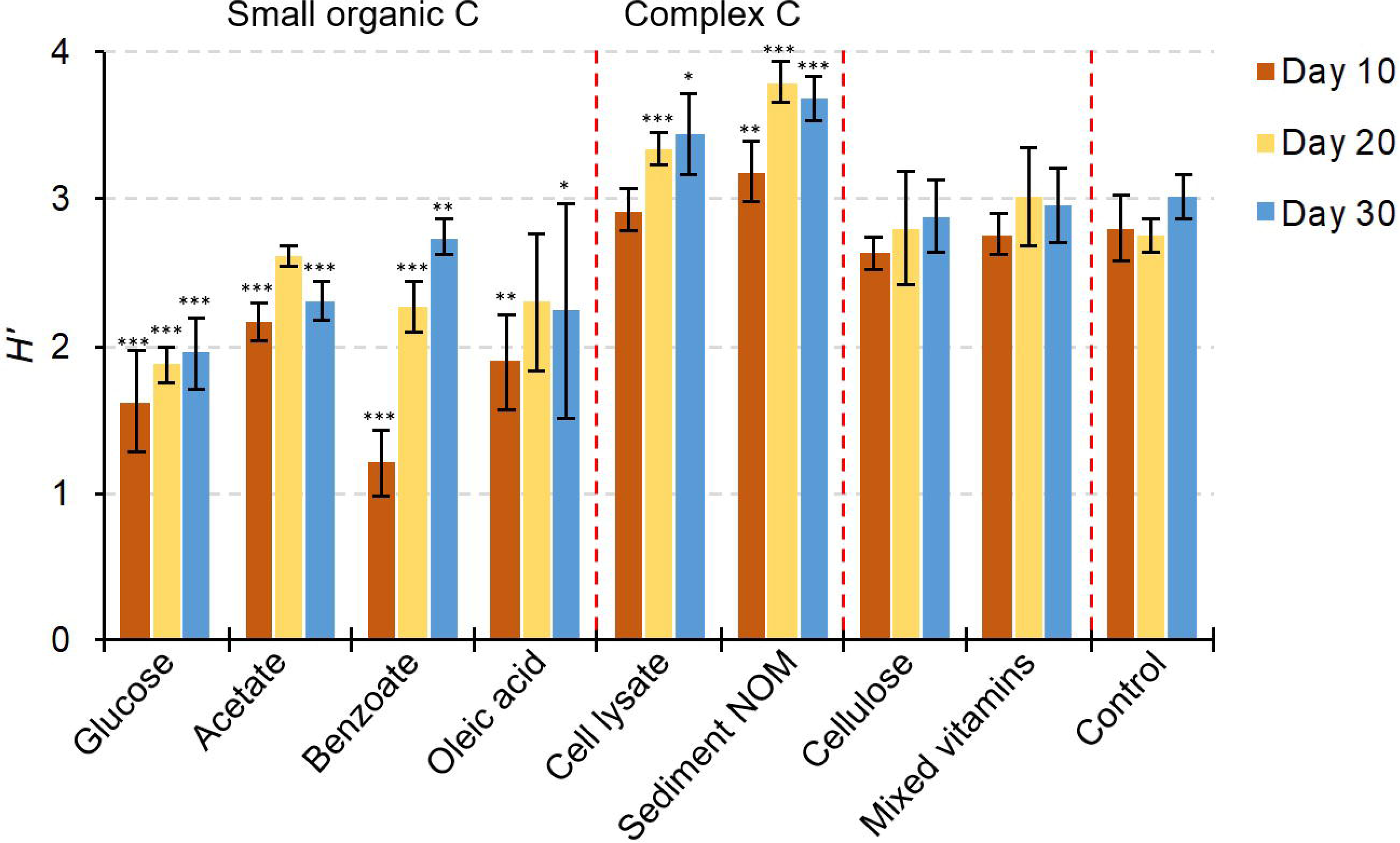
Bacterial diversity of enrichments amended with different C sources, as indicated by Shannon’s diversity index (*H’*). Significance between experimental group and corresponding control group is indicated by *** when *p* < 0.001, ** when *p* < 0.01, and * when *p* < 0.05.

### Complex C sources enriched rarely cultivated bacterial taxa

We further studied taxonomic responses of subsurface groundwater bacterial communities to different C sources. Out of the quality-filtered reads, 21 phyla and 94 orders were taxonomically identified, covering 71–100% of entire reads, except two samples (57% and 60%) in bacterial cell lysate-amended group. All phyla and 34 abundant orders (with relative abundance >1% in any sample) are presented in Figure 3 and Figure 4, respectively.

**Figure 3.**
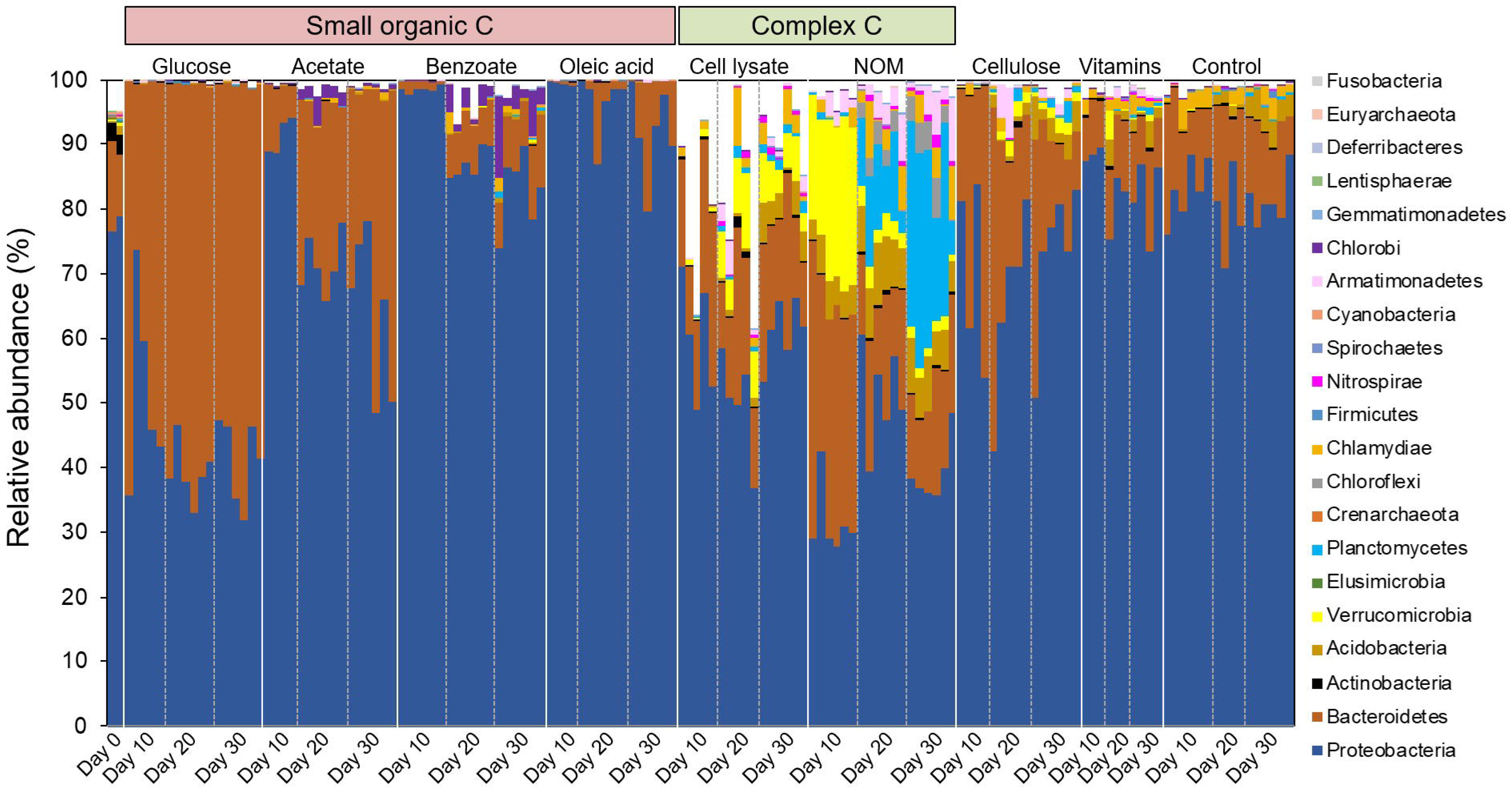
Relative abundance of each taxonomic phylum.

**Figure 4.**
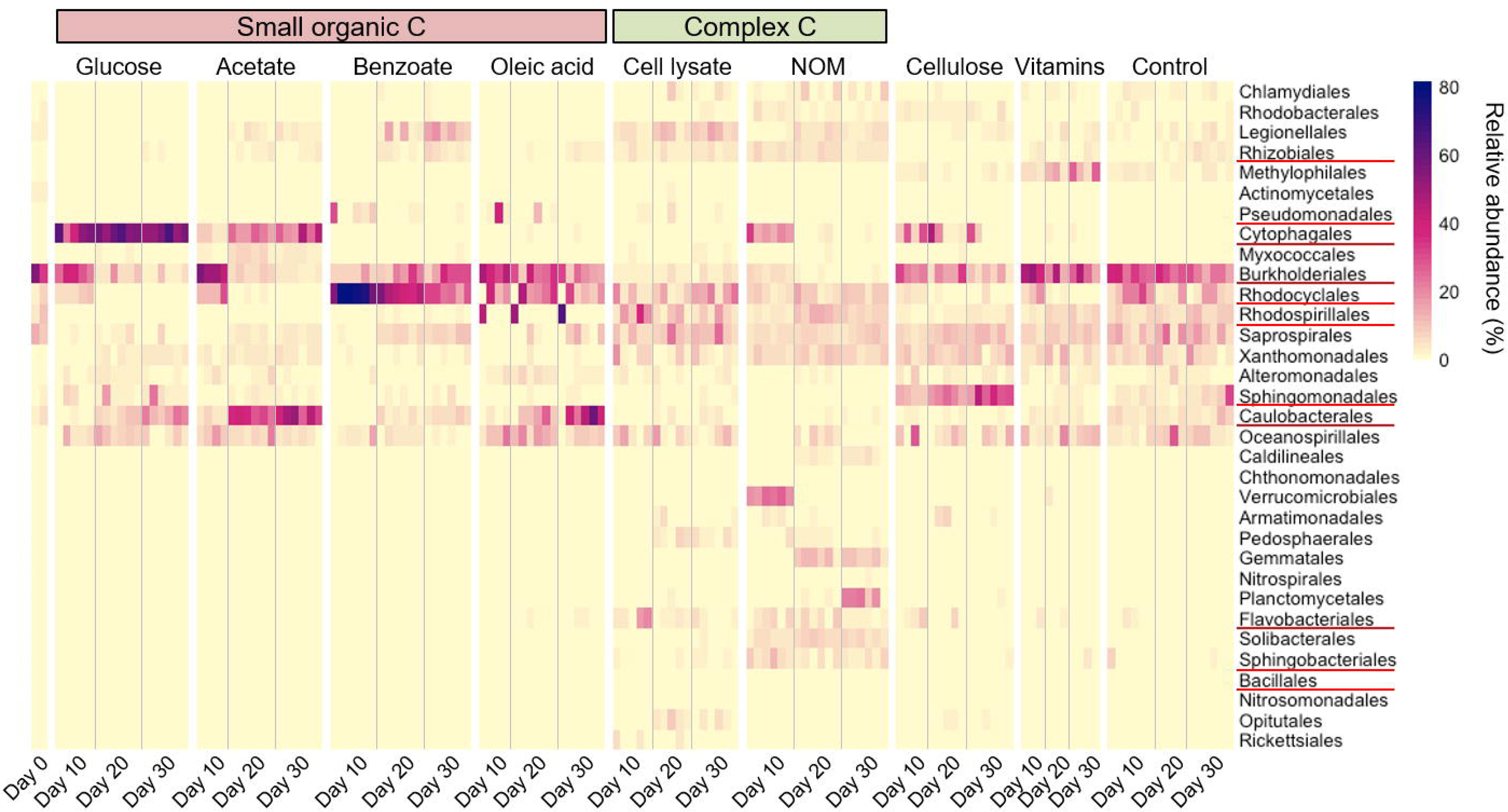
Relative abundance of each taxonomic order (> 1% in any sample). Orders having representative isolates in this study were marked with red underlines.

Except for glucose-amended enrichments, *Proteobacteria* was the most dominant phylum especially in benzoate- and oleic acid-amended groups (Figure 3). *Bacteroidetes* was mostly dominant over *Proteobacteria* in glucose-amended enrichments, and was also highly abundant in other groups (Figure 3). It is worth noting that *Verrucomicrobia* and *Planctomycetes* were present abundantly in cultures with sediment NOM as the C source. Only a handful isolates of *Verrucomicrobia* have been successfully cultivated thus far^5, 33, 34^, although members of this bacterial phylum are ubiquitous in the environment^35, 36^. *Planctomycetes* is of deep interest to microbiologist due to their unique and peculiar characteristics, and only ~2% of strains in this phylum have been isolated in pure culture^37^. These two phyla existed in NOM-amended enrichments with a clear succession pattern. *Verrucomicrobia* was highly abundant at an early stage and significantly diminished over time, while *Planctomycetes* became one of major phyla at late stages (Figure 3). It was reported that *Planctomycetes* are comparatively slow growing organisms with low demand for C and nitrogen sources^37^, which may explain their late appearance in the cultures. The phylum *Armatimonadetes* was also abundantly present in NOM-amended enrichments. *Armatimonadetes* has long been lacking of isolated member until 2011^38^, and so far only 3 isolated strains in this phylum have been reported^38-40^.

At the order level, microcosms amended with simple small organic C highly enriched a few orders such as *Cytophagales, Burkholderiales, Rhodocyclales, Caulobacterales*, and *Oceanospirillales* (Figure 4). As a comparison, in complex C-amended microcosms, diverse orders were enriched, including those that were hardly enriched in other groups, e.g., *Verrucomicrobiales, Gemmatales, Planctomycetales, Flavobacteriales, Solibacterales*, and *Sphingobacteriales* (Figure 4).

### C utilization pattern

Different bacterial species may have different preferable C sources when growing in culture medium under laboratory condition. In this study, species that were enriched on at least one C source were selected and classified as generalist, intermediate, or specialist based on the criteria described above. As shown in Figure 5, the 4 generalists (that can be enriched on all types of C sources in this study) include one *Betaproteobacteria* (*Pelomonas* sp.), two *Gammaproteobacteria* (*Halomonas* sp. and *Shewanella* sp.), and one *Bacteroidetes* (*Sediminibacterium* sp.), indicating that these species likely harbor the metabolic potential of utilizing diverse C sources, from simple small organic C to undefined complex C. There are 30 intermediates that were enriched on 2–8 types of C sources, distributing in *Alpha*-, *Beta*-, *Gamma*-*proteobacteria, Bacteroidetes, Chlamydiae*, and *Verrucomicrobia* (Figure 5).

**Figure 5.**
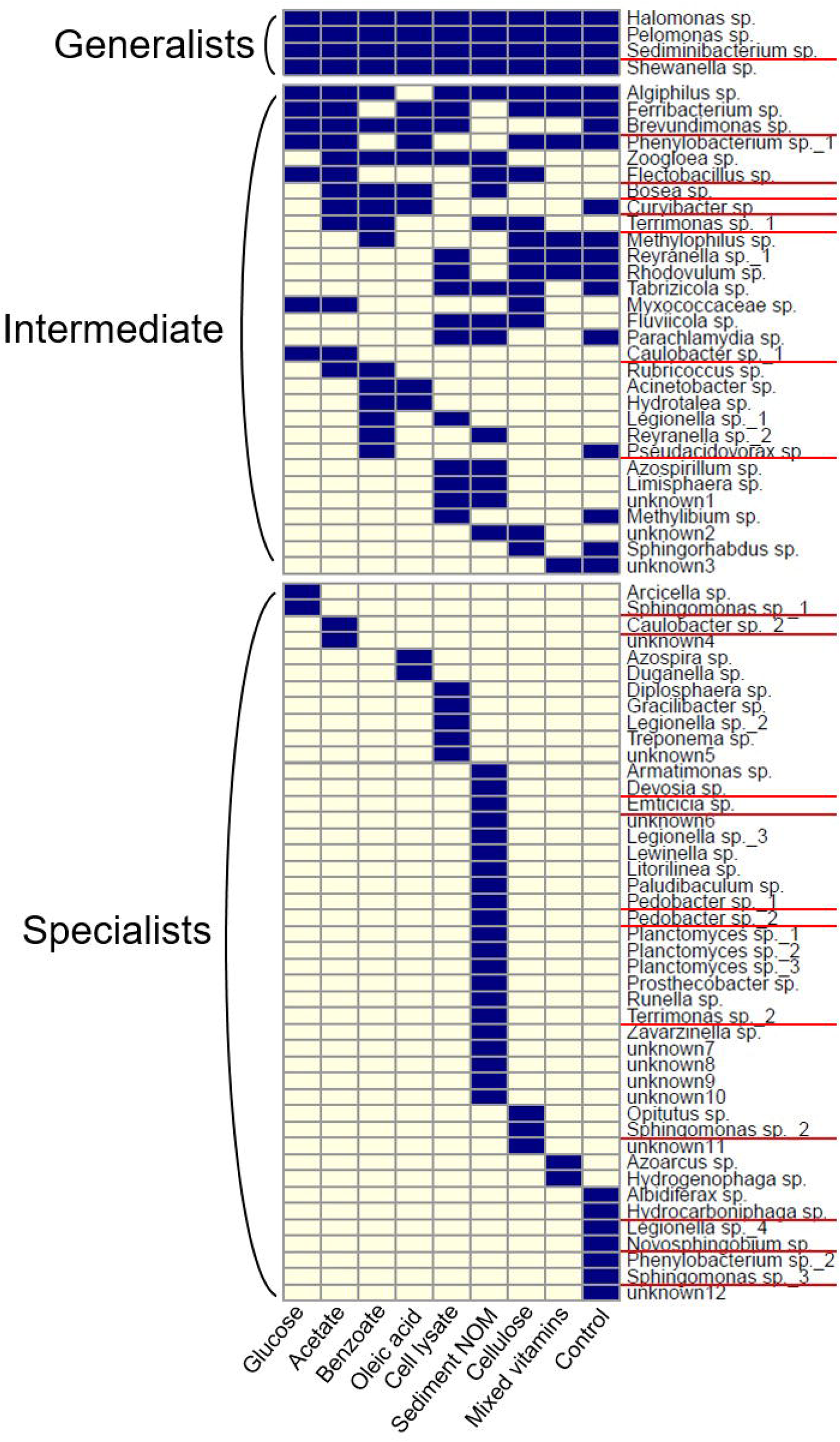
Enriched bacterial species (>1% in at least three replicates for any time point) on different C sources. Species having representative isolates in this study were marked with red underlines.

A total of 44 specialists that were exclusively enriched on specific type of C source are identified, a half of which were preferably grown with sediment NOM, including some novel (unclassified) species and those are rarely cultivated in the lab such as *Armatinomas* sp. and *Planctomyces* sp. (Figure 5). Some species within the same genus differed in C utilization pattern. For example, *Legionella* sp._1 could be enriched on benzoate and bacterial cell lysate, while *Legionella* sp._2, 3, and 4 were only enriched on bacterial cell lysate, sediment NOM, and groundwater indigenous NOM (control group), respectively (Figure 5).

### Isolates from complex C-amended microcosm enrichments

From the enrichments amended with complex C (bacterial cell lysate or NOM), we were able to isolate and cultivate a total of 271 pure strains of bacteria, including fast-growing (within 2 days) and slow-growing (up to 3 weeks) isolates. These isolates were grouped into 51 species, representing 4 phyla and 13 orders (Table 1).

Based on 16S rRNA similarity to published species, candidates of novel species were defined by comparison of 16S rRNA similarity at a threshold of 98%, novel genus level at 90–95%, and novel family level at an off-limit lower than 90%^22^. According to this criteria, five novel bacteria were isolated from subsurface groundwater in this study, including three novel species: FW305-C-84 (*Dyadobacter sediminis* strain Z12, 97%), FW305-C-176I (*Rhizobium rosettiformans* strain W3, 96%), and FW305-C-3 (*Sinimarinibacterium flocculans* strain NH6-24, 96%); one novel genus candidate: FW305-C-2 (*Panacagrimonas perspica* strain Gsoil 142, 93%); and one novel family candidate: FW305-C-21 (*Solitalea canadensis* strain DSM 3403, 86%).

The isolation results were then compared with 16S rRNA gene surveys of enrichments in order to see how efficient complex C can serve as a substrate for diverse bacterial isolation. Of high significance is the fact that we were successful in cultivating one-third (11 out of 33) of the enriched orders identified by the molecular technique through isolation efforts (Figure 4). Matched isolates were also obtained for a quarter (19 out of 78) of enriched species detected by the molecular method, including 1 generalist, 7 intermediates, and 11 specialists (Figure 5).

## Discussion

An often-invoked principle in microbial ecology is the notion that “everything is everywhere, and the environment selects.” Under this paradigm, we expect that subsurface bacteria communities would respond differently to different C sources in culture medium. In this study, we carried out microcosm experiments in the lab and provided intrinsic bacterial communities from freshly collected groundwater amended with different C sources, including simple small organic C (glucose, acetate, benzoate, or oleic acid), undefined complex C (bacterial cell lysate or NOM), polysaccharide (cellulose), and vitamins mix. Results showed that greater diversity of bacteria was recovered from subsurface groundwater under laboratory cultivation condition by using complex C sources. Some rarely cultured phyla such as *Verrucomicrobia, Planctomycetes*, and *Armatimonadetes* were enriched on sediment NOM, but not on traditional simple small organic C sources (Figure 3). The small organic C only enriched a few phyla which are commonly isolated and already have diverse representative isolates (Figure 2). As the major naturally occurring C sources for microbes in the subsurface, sediment NOM are mixtures of heterogeneous organic substrates containing proteins, nucleic acids, lipids, carbohydrates, etc^41^. These undefined complex C support various subsurface bacteria in culture medium under laboratory condition, which may benefit for cultivation and isolation of key bacteria species from subsurface environment.

Further isolation led to 51 species from four different phyla, including 5 novel isolates by using complex C-amended enrichments as inoculums on pour agar plates (Table 1). Most isolates belonged to the most dominant phylum *Proteobacteria*. Although being enriched in liquid cultures, species from phyla *Verrucomicrobia, Planctomycetes*, and *Armatimonadetes* failed to grow as pure colonies on agar plates in this study, suggesting that other isolation techniques such as serial dilution are needed in order to get pure cultures of these bacteria. In total, about one-third of enriched orders and one-fourth of enriched species known from molecular surveys were recovered by isolation and had pure cultured representatives (Figure 4 and 5). The recovery of multiple species is encouraging and indicates progress toward a better recovery of diverse microbes from the subsurface.

Cultivation attempts undertaken at several FRC sites have produced limited numbers of microbial isolates^9, 42, 43^. Fields et al.^43^ used conventional direct plating method with nitrate amended nutrient broth or MR2A medium for bacterial isolation from one FRC groundwater (background) sample, obtaining 13 species classified in two phyla: *Proteobacteria* (*Alpha-, Beta-, Gamma-*, and *Delta-*) and *Actinobacteria*. Bollmann et al.^9^ applied *in situ* diffusion chamber incubation technique to isolate bacterial species from FRC contaminated sediment. They obtained 61 strains and 50 species representing *Proteobacteria* (*Alpha*-, *Beta*-, and *Gamma*-), *Bacteroidetes, Actinobacteria*, and *Firmicutes*, the same phyla as we obtained in this study. They also used conventional direct plating method for isolation and obtained only 8 species representing *Proteobacteria* (*Alpha-* and *Gamma-*), *Verrucomicrobia*, and *Actinobacteria*. Compared to diffusion chamber-based approach, the method used in this study is an easier application in laboratories to obtain comparably diverse bacterial isolates.

In summary, this study shows natural complex C substrates may enrich much more diverse bacterial communities from subsurface groundwater compared to traditional simple small organic C sources. Additionally, isolation was simpler and could be directly subcultured to obtain bacterial pure cultures for further investigation of physiology and geochemical roles of key species in subsurface ecosystems.

## Acknowledgement

This material by ENIGMA-Ecosystems and Networks Integrated with Genes and Molecular Assemblies (http://enigma.lbl.gov), a Scientific Focus Area Program at Lawrence Berkeley National Laboratory is based upon work supported by the U.S. Department of Energy, Office of Science, Office of Biological & Environmental Research under contract number DE-AC02-05CH11231. The FRC groundwater sample was kindly provided by Terry C Hazen and Dominique C Joyner from Oak Ridge National Laboratory. We would also like to thank the MIT BioMicro Center for sequencing support. The sequencing efforts were funded by the National Institute of Environmental Health Sciences of the NIH under award P30-ES002109.

