## Supplementary for "Capturing the Diversity of Subsurface Microbiota – Choice of Carbon Source for Microcosm Enrichment and Isolation of Groundwater Bacteria"

**Supplementary Information**

Ingredients of stock solution of vitamins.

| **Chemical** | **Concentration (mg/L)** |
| --- | --- |
| Vitamin B_1_ (thiamine) | 500 |
| Vitamin B_2_ (riboflavin) | 500 |
| Vitamin B_3_ (nicotinic acid) | 500 |
| Vitamin B_5_  (pantothenic acid) | 500 |
| Vitamin B_6_ (pyridoxine HCl) | 1000 |
| Vitamin B_7_ (biotin) | 200 |
| Vitamin B_9_ (folic acid) | 200 |
| Vitamin B_10_ (p-amino benzoic acid) | 500 |
| Vitamin B_12_ (cobalamin) | 10 |
| D,L-6,8-thioctic acid | 500 |

Table S1. Primers used for 16S rRNA gene amplicon sequencing.

| **Primer name** | **Primer sequence (5’ -> 3’)** |
| --- | --- |
| PE16S_V4_U515_F* | ACACGACGCTCTTCCGATCTYRYRGTGCCAGCMGCCGCGGTAA |
| PE16S_V4_E786R* | CGGCATTCCTGCTGAACCGCTCTTCCGATCTGGACTACHVGGGTWTCTAAT |
| PE-III-PCR-F-### | AATGATACGGCGACCACCGAGATCTACACNNNNNNNNACACTCTTTCCCTACACGACGCTCTTCCGATCT |
| PE-IV-PCR-R-### | CAAGCAGAAGACGGCATACGAGATNNNNNNNNCGGTCTCGGCATTCCTGCTGAACCGCTCTTCCGATCT |

*Universal primer segments U515 and E786R were drawn from (Lane, 1991).
